# Predicting and phase targeting brain oscillations in real-time

**DOI:** 10.1101/2024.10.12.617974

**Authors:** Stefan L. Jongejan, Anna C. van der Heijden, João Patriota, Lucia M. Talamini

## Abstract

**Objective:** Closed-loop neurostimulation (CLNS) procedures, aligning stimuli with electrical brain activity, are quickly gaining popularity in neuroscience. They have been employed to reveal causal links between neural activity patterns and function, and to explore therapeutic effects of electroencephalography (EEG-)guided stimulations during sleep. Most CLNS procedures are developed for a single purpose, detecting one specific pattern of interest in the EEG. Furthermore, most procedures have limited, if any, flexibility to adapt to temporal or interindividual variance in the signal, which means they wouldn’t work optimally across the full physiological phenomenology.

**Approach:** Here we present a new approach to CLNS, based on real-time signal modelling to predict brain activity, allowing targeting of a broad variety of oscillatory dynamics. Intrinsic to the modelling approach is adaptation to signal variance, such that no personalization steps prior to use are necessary. We systematically assess stimulus targeting performance of modelling-based CLNS (M-CLNS), across a wide range of brain oscillation frequencies and phases in human and rodent neurophysiological signals.

**Main results:** Our results show high performance for all target phases and frequency bands, including slow oscillations, theta and alpha waves.

**Significance:** These findings highlight the general applicability and adaptability of M-CLNS, which also favors its application in populations with an atypical oscillatory signature, like clinical or elderly populations. In conclusion, M-CLNS provides a promising new tool for neural activity-dependent stimulation in both experimental research and therapeutic applications, such as enhancing deep sleep in patients with sleep disorders.

## 1. Introduction

Recent advances in neurotechnology have led to the development of automated procedures for aligning stimuli with electrical brain activity. Such neurally-guided stimulation methods entail the continuous, real-time, processing of short stretches of signal to detect a particular signal characteristic, upon which a stimulus is sent out to interact with neural processing. In reported studies, the neurophysiological signal has ranged from electroencephalography (EEG) [1] to local field potentials (LFP) [2,3], to the action potential patterns of individual neurons [4]. Stimuli have included sensory (e.g. auditory or visual) and electrical ones [3,5-7]. Such closed-loop neurostimulation (CLNS) methods have opened up new avenues in experimentation, revealing direct and causal links between different neural activity patterns and their function [8-12]. Moreover, CLNS methods have high clinical relevance, with potential applications in many types of therapeutic stimulation, including deep brain or nerve stimulation and non-invasive sleep enhancement [13-16].

Much of the effort regarding CLNS stems from the field of sleep, where CLNS has offered a means to circumvent the experimental challenges posed by the paucity of state-related behavior. CLNS has, for instance, been used to target stimuli to relatively long-lasting brain-body states, such as rapid eye movement (REM) or non-rapid eye movement (NREM) sleep [17,18]. More challenging applications aimed to align stimuli with fleeting dynamics, such as oscillations in a particular frequency band, or even a particular phase of such oscillations. Examples are the interruption of sleep spindles or sharp wave ripples with electrical stimuli, during NREM sleep, demonstrating their causal involvement in memory consolidation [19,20]. Other examples regard targeting the presentation of auditory memory cues to (a particular phase of) slow oscillations, another NREM sleep dynamic, to influence memory reprocessing [21]. Both types of experiment have contributed importantly to elucidating the causal role of sleep-related oscillatory dynamics in information processing.

In most of the aforementioned studies, the CLNS procedure was dedicated to targeting one specific pattern in the physiological signal. In such cases, pattern detection is typically based on signal filtering in the frequency range of interest and application of threshold criteria on the envelope of the filtered signal (e.g. amplitude threshold crossings and duration of such crossings). These ‘dedicated’ procedures are not adaptive to temporal or interindividual variance in the signal. Furthermore, stimulus presentation necessarily lags after pattern detection. These characteristics, at least on theoretical grounds, would limit precision in targeting any highly time variant dynamic. Also, given the lack of adaptability, such methods typically require individualization by adjusting the CLNS algorithm’s threshold criteria during a so-called baseline night. The use of threshold criteria, including amplitude thresholds may result particularly difficult in (clinical) populations in which the target dynamic is weakened or otherwise deviates from the norm.

Another putative approach to detecting a specific pattern of interest in an ongoing signal is through artificial neural networks. These, however, need to be trained on a large data set and tend to have relatively long execution times, which may hinder real-time applications [22].

Still another approach to CLNS, involves phase-locked loops (PLL). Specifically, it has been suggested that a PLL could track the phase of slow oscillations, which could be used to release a stimulus upon detection of a given phase [23-25]. A PLL matches the phase of a single defined sinusoidal curve to a signal, and only detects zero crossings. Given these characteristics, it is unlikely that a PLL could accurately track the phase of any brain-electrical signals, which are highly time-variant and do not feature dominant frequencies that stick out sharply in the frequency spectrum. A higher level of sophistication, compared to previously mentioned approaches, can be achieved using a modelling approach to CLNS. This holds that the signal is modelled in real-time, allowing for prediction of the ongoing signal through model extrapolation. A modelling approach entails adaptability to the signal and, moreover, allows stimuli to be targeted to events detected in the prospected signal, rather than lagging after the detected events.

A first modelling-based CLNS approach, developed by our group, used a sine fitting procedure on data filtered in a 1 Hz range around the momentary slow oscillation center frequency, followed by a Hilbert transform [26]. Extrapolation of the model allowed sending out stimuli to coincide with a target phase detected in the forecasted signal. This method was shown to perform reasonably well in targeting stimuli to slow oscillation phases.

As indicated previously, most of the aforementioned procedures were developed to target one specific pattern in a physiological signal. Oscillatory phase-targeting (21-26), arguably one of the most challenging aims for closed-loop procedures, has to our knowledge only been reported for EEG slow waves, a slow dynamic (~0.8 Hz) characteristic of deep sleep. Phase-targeting performance, when reported, has been varying and no method has been thoroughly tested for its performance across a wide range of phase angles. Indeed, existing CLNS methods have been limited to targeting narrow ranges of frequencies, phases, or activities.

We here present a new, modelling-based, CLNS method (M-CLNS) that allows flexible targeting of different signal frequencies and phases and has been utilized to precisely phase-lock stimuli to brain oscillations during various sleep-wake stages [27-29]. The method, based on non-linear sine fitting of the raw neurophysiological signal, is fully adaptive and can, in principle, be applied to any signal with oscillatory characteristics. Model extrapolation delivers a near-continuous stream of predictions of the oscillatory progression of the future signal, irrespective of the momentary signal content. Thus, any phase of any oscillatory frequency can, in principle, be targeted, as can any other pattern detected in the prospected activity. The real-time algorithm operates on short data stretches, which, together with its short processing time (~2 ms using any up-to-date processor), favors application to highly time variant dynamics and high oscillatory frequencies. The omission of fast filtering, used in other procedures also favors speed and avoids signal distortion, which can negatively affect targeting performance. Due to the method’s intrinsic adaptability to the signal, M-CLNS can, in principle, be directly applied to any subject; no algorithm training or adaptation night for calibration of the algorithm settings are necessary, saving time and resources.

While real-time modelling and model extrapolation in M-CLNS occur continuously, stimulus release is dictated by the following criteria: (1) the frequency of the model is within a predefined range of interest; (2) the normalized cross correlation coefficient between the model and raw signal segment is above a predefined minimal value; (3) the targeted phase is predicted to occur in the forecast signal within a predefined future time interval, the so-called search window. This search window is set to account for hardware lag and analysis time; it extends only a short time into the future in view of the high time variance of electrophysiological neural signals. Optionally, additional criteria can be set, such as a minimum inter-stimulation interval, or a minimum amplitude threshold of the signal in the fitting segment. If all set criteria are met, a stimulus is released to coincide with the upcoming predicted target phase, accounting for loop lag (processing time of the algorithm plus the throughput time of all other hardware and software components in the loop) and the future time point of the predicted phase.

Here, we systematically assess phase targeting performance of M-CLNS, across a wide range of oscillatory dynamics and phases, in both human and rodent neurophysiological signals. Specifically, we target hallmark oscillations of deep sleep (slow oscillations, 0.5-1.5 Hz), REM sleep (theta waves, 4-8 Hz) and waking (alpha waves, 8-12 Hz), using concatenated EEG data for humans and hippocampal LFP data for rodents. We use a simulation approach to systematically assess targeting performance for eight different target phases equidistantly spread across the 360° range, for each hallmark oscillation. For each simulation, we provide measures of phase targeting performance, including accuracy, precision, and stimulation density. In addition, to evaluate the consistency of performance across different phases and participants, we calculate performance for each targeted phase (merged across participants) and for each participant (merged across targeted phases), for each hallmark oscillation.

## 2. Methods

### 2.1. Human polysomnography

Human sleep-EEG data was obtained from pre-existing high-density polysomnography (PSG) recordings that were previously collected during nocturnal sleep studies at our lab. These recordings were made in healthy human participants that were given an at least 8-hour sleep opportunity at the sleep lab and were not exposed to any manipulations either during the PSG night or during previous nights. PSG recordings were obtained using sized WaveGuard caps (ANT, Enschede, The Netherlands) with separate sintered reference electrodes on the earlobes (A1, A2). The EEG signals, together with bipolar vertical- and horizontal-EOG and chin EMG signals, were captured using a Refa 8, 72-channel, DC amplifier (TMS international, Oldenzaal, The Netherlands), with a sampling rate of 512 Hz. The acquired signals were recorded on a dedicated computer using TMSi Polybench (1.34.1.4456). EEG impedances were confirmed to be below 10 kΩ at the start of each recording.

The PSG recordings were sleep stage scored according to AASM guidelines [30]. Hereafter, three data sets were created based on discrete sleep stages. Each data set contained the Fpz-A1A2 EEG signal, which was high-pass filtered at 0.1 Hz to attenuate the DC component: (1) a slow oscillations data set, consisting of 140 minutes of deep sleep data obtained from only N3 sleep of seven participants (4 males; age M = 21.14, SD = 1.21); (2) a theta waves data set comprising 120 minutes of only REM sleep obtained from six other participants (4 males; age M = 21.67, SD = 2.80); (3) an alpha waves data set, consisting of 84 minutes of only wake after sleep onset (WASO) data without body movements, obtained from five more participants (1 male; age M = 20.60, SD = 1.14). All three data sets were used as input to the M-CLNS algorithm, to determine M-CLNS’ targeting performance for each hallmark oscillation.

### 2.2. Tetrode local field potentials in rodents

Rodent LFP sleep data was obtained through tetrode recordings from the dorsal hippocampal CA1 area (target coordinates in mm: −3.2 AP, −1.51 ML, 2.88 DV) in three male Lister Hooded rats (aged 2-5 months; Envigo, The Netherlands). These animals were used in a related research project at our lab. Tetrodes were constructed from four twisted 13-µm coated nichrome wires (California Fine Wire, Grover Beach, United States) with gold-plated tips to reduce electrode impedances. The tetrodes were connected to an Open Ephys acquisition board (www.open-ephys.org) through operational amplifiers (RHD2000, Intan) and tether cable, connected to a Saturn-5 commutator (Neuralynx, Bozeman, United States). The data was sampled at 30 kHz, recorded on a dedicated computer, and resampled to 512 Hz in EventIDE. Animals were equipped with a headstage (Intan RHD2132 board) that included a 3-axis accelerometer, giving access to the animals’ acceleration with high temporal resolution, allowing for sleep stage scoring. A data set for simulations was created consisting of 24.25 minutes of theta waves during REM sleep from the three rats. This data set was used as input to the M-CLNS algorithm, to determine the M-CLNS’ targeting performance.

### 2.3. M-CLNS algorithm

The M-CLNS algorithm used in this experiment was developed at our lab in collaboration with Okazolab Ltd, London, UK, and was incorporated in a custom version of the visual programming software EventIDE (v.25.4.2022). The M-CLNS algorithm continuously performs non-linear sine fitting on the most recent segment of the raw incoming signal, the calculus window. The best-fitting estimated model is used to extrapolate the signal into the future and the first occurrence of a pre-defined oscillatory target phase is found in the forecasted signal. As mentioned in the introduction, for stimulus release to occur at the identified target phase, criteria regarding the model frequency, normalized cross correlation coefficient between model and raw signal segment, and search window need to be met. The minimum search window time used for the current study was 10 ms. The minimum time between consecutive stimulus presentations used for this study was 500 ms. Parameter settings specific to the targeted frequency are listed in the next section.

### 2.4. Simulations

The four data sets, comprising three human EEG recordings and one rodent LFP recording, were used for offline simulations in EventIDE. During simulations, the raw online brain signals were emulated, to test the M-CLNS algorithm performance. Each data set was used for eight simulations, to assess performance on eight different target phases. The targeted phases were 0°, 45°, 90°, 135°, 180°, 225°, 270°, and 315° (figure 1i). The raw signal together with the stimulus presentation onset markers were recorded to an output file during each simulation.

**Figure 1.**
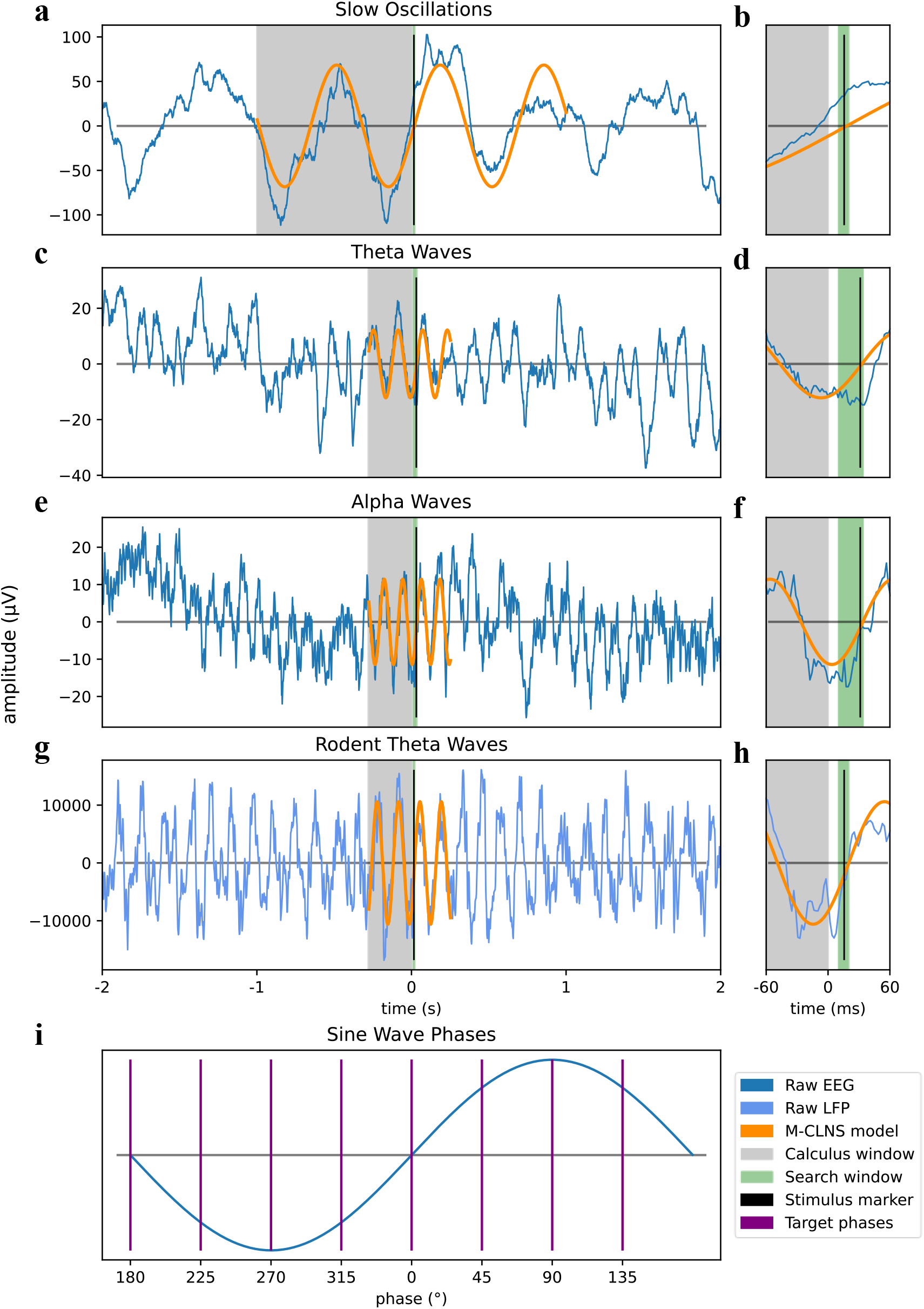
M-CLNS modelling of different brain oscillations. (a-h) Examples of the M-CLNS modelling, together with an enhanced view of the future search window and stimulus marker, are shown for human EEG slow oscillations (a,b), theta waves (c,d), alpha waves (e,f) and rodent LFP theta waves (g,h). Each graph shows raw EEG or LFP signals (blue) with the M-CLNS model (orange), based on the calculus window (grey). The future search window (green) contains the recorded stimulus presentation moment (black). (i) Schematic graph showing the location of the eight targeted phases (0°, 45°, 90°, 135°, 180°, 225°, 270°, and 315°) as part of a sinewave.

The deep sleep data set was used to target slow oscillations using the most recent 1000 ms (512 samples) in the calculus window, with a minimum normalized cross correlation coefficient of 0.3 and a maximum search window time of 20 ms. The REM sleep data set was used to target theta waves using the most recent 280 ms (143 samples) with a minimum normalized cross correlation coefficient of 0.7 and a maximum search window time of 20 ms. In addition, a minimum signal amplitude threshold of 8 µV during the calculus window was set. The WASO data set was used to target alpha waves using the most recent 280 ms (143 samples) with a minimum normalized cross correlation coefficient of 0.3 and a maximum search window time of 34 ms. The rodent REM sleep data set was used to target theta waves using the most recent 280 ms (143 samples) of the raw LFP signal. The minimum normalized cross correlation coefficient was 0.8, and the maximum search window time was 20 ms.

Of note, the algorithm parameter settings adopted for targeting different frequency bands, were set based on theoretical notions (e.g. the calculus window on which real-time analyses were done is shorter for the faster dynamics) and further fine-tuned through simulations. However, no exhaustive optimization procedures were performed.

### 2.5. Data analysis

Data analyses were performed in Python (3.10.121) using the Pandas (2.1.4), NumPy s(1.26.2), MNE (1.6.0), SciPy (1.11.4), PyCircStat (0.0.2), Astropy (6.0.0), and Matplotlib (3.8.2) libraries [31-38]. The output files, containing the raw EEG and LFP signals along with stimulus onset markers, were converted to MNE raw objects and bandpass filtered using a finite impulse response (FIR) filter with frequency ranges matching those targeted by the M-CLSN algorithm (SO: 0.5-1.5 Hz, theta waves: 4-8 Hz, alpha waves: 8-12 Hz).

For each of the 32 filtered output data sets, the analytic signal was obtained by applying a Hilbert transform, and the instantaneous phase was derived at each stimulus marker. To correct for the phase shift induced by the Hilbert transform, a correction was applied by subtracting π/2 from each instantaneous phase. The stimulus marker offset from the targeted phase, in degrees, was then calculated for each corrected instantaneous phase, performed separately for each of the eight targeted phases.

Circular statistics, including the mean, median and standard deviation of the phase offsets, were computed for each of the 32 output data sets to determine the accuracy and precision of stimulus presentations. Smaller mean and median phase offsets signify higher accuracy of stimulus presentations. Smaller phase offset standard deviations signify higher precision of stimulus presentations. These statistics were also calculated across all eight targeted phases corresponding to the same hallmark oscillation (grand averages), as well as for each participant across all eight targeted phases of the same hallmark oscillation.

To assess stimulation density, the inter-stimulus-interval between consecutive stimulus onset markers was computed. Average inter-stimulus-interval values were then calculated per targeted phase of each hallmark oscillation, per participant across all target phases of each hallmark oscillation, and collapsed over target phases and participants per hallmark oscillation (grand average). Lastly, as a secondary measure of stimulus spread, we calculated the percentage of stimuli presented within a 180° range centered around the target phase and determined the same averages as for the other performance measures.

## 3. Results

### 3.1. M-CLNS adequately models varied oscillatory dynamics in different sleep-wake states

M-CLNS was able to adequately track and predict all hall mark oscillations, that is, slow oscillations during deep sleep, theta waves during REM sleep and alpha waves during wake in human EEG recordings, as well as REM sleep theta in hippocampal LFP recordings in rodents. Representative examples of M-CLNS modelling and prediction for each hall mark oscillation are shown in figure 1.

### 3.2. Slow oscillations during deep sleep in human EEG

For slow oscillations in human EEG, the grand average performance, expressing the offset from the target phase across all eight targeted phases and all participants, was 1.85° ± 49.16° (mean ± SD), based on 36693 stimulus onset markers (see figure 2a). Of all markers, 87.32% were recorded within a 180° range centered around the target phase. The inter-stimulus-interval averaged across all markers was 1.82 ± 1.59 seconds (mean ± SD). Given that slow oscillations have a maximum duration of about two seconds (0.5 Hz), this suggests that suggests that most naturally occurring slow oscillations were targeted.

**Figure 2.**
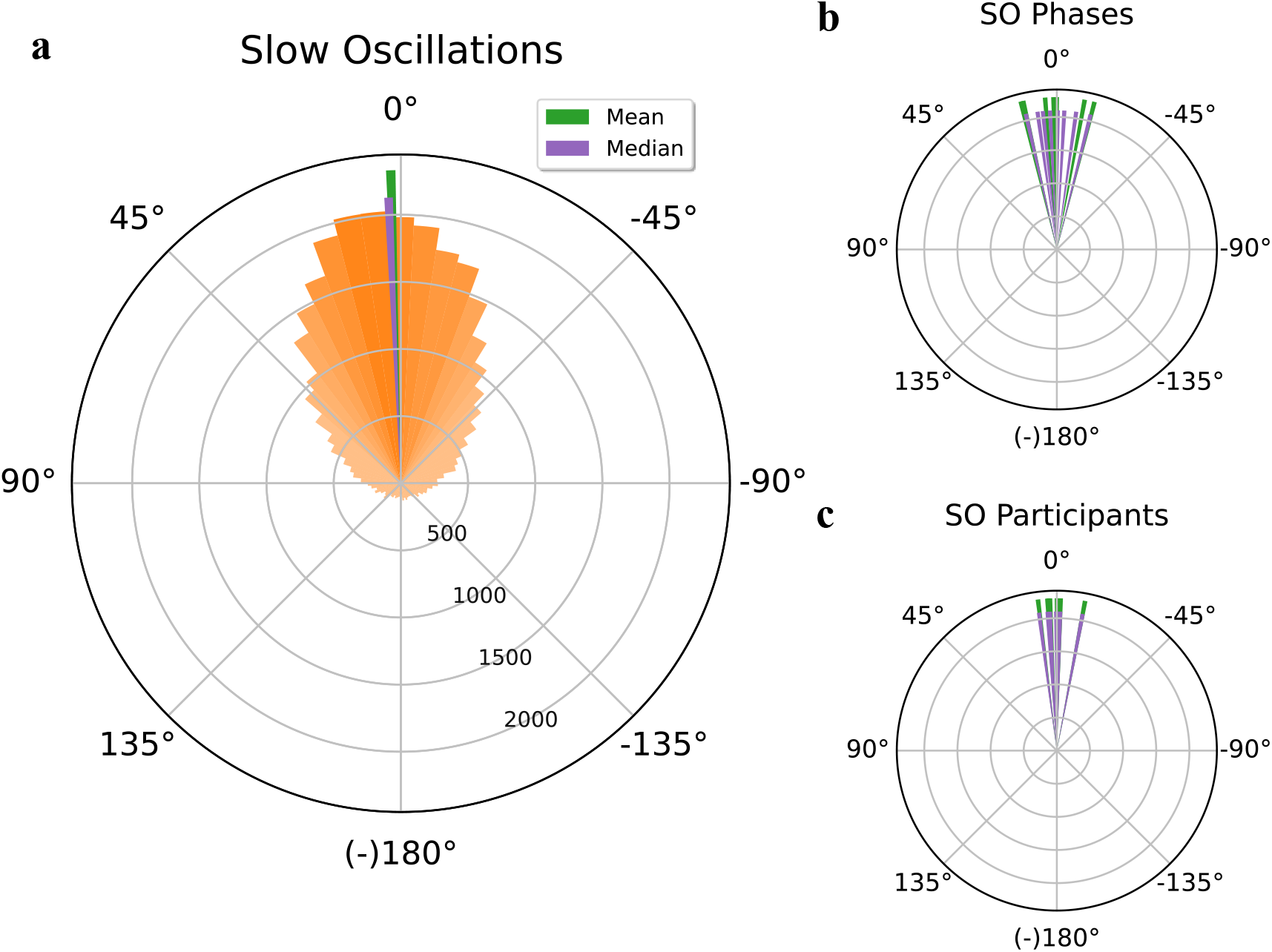
Phase-targeting performance for slow oscillations in human EEG. (a) Circular histogram showing the distribution of binned stimulus marker phase offsets, defined as the difference between targeted phase and the actual stimulus marker phase, measured in degrees. The circular histogram combines data from all eight targeted phases (0°, 45°, 90°, 135°, 180°, 225°, 270°, and 315°) and all participants (orange) together with the mean (green line) and median (purple line) stimulus marker phase offset. (b) Circular histogram depicting the mean (green lines) and median (purple lines) stimulus marker phase offset for each of the eight individual targeted phases. (c) Circular histogram depicting the mean (green lines) and median (purple lines) stimulus marker phase offset for each individual participant.

Accuracy and precision were high for all individual target phases (table 1 and figure 2b; see supplementary figure 1 for polar plots per targeted phase), ranging between a phase offset of 0.05° ± 46.09° (mean ± SD) and −14.32° ± 46.45°. The inter-stimulus-interval ranged between 1.53 ± 1.14 and 2.18 ± 2.02 seconds.

**Table 1.** Phase-targeting performance for individual and across combined targeted phases (0°, 45°, 90°, 135°, 180°, 225°, 270°, and 315°), for slow oscillations, theta and alpha waves in human EEG.

| Target phase | Phase offset (°) |  |  | 180° range (%) | Inter-stimulus-interval (s) |  |
| --- | --- | --- | --- | --- | --- | --- |
|  | Mean | SD | Median | Percentage | Mean | SD |
| **Slow oscillations in human EEG** |  |  |  |  |  |  |
| **0** | 13.71 | 49.09 | 12.9 | 84.64 | 1.52 | 1.14 |
| **45** | 12.78 | 43.61 | 7.99 | 89.59 | 1.67 | 1.34 |
| **90** | 0.05 | 46.09 | -3.09 | 90.84 | 2.03 | 1.93 |
| **135** | -14.32 | 46.45 | -14.05 | 89.61 | 2.18 | 2.02 |
| **180** | -10.55 | 52.34 | -8.02 | 84.07 | 1.65 | 1.38 |
| **225** | 4.43 | 48.72 | 2.82 | 87.65 | 1.75 | 1.44 |
| **270** | 0.47 | 51.19 | -0.37 | 86.97 | 1.96 | 1.69 |
| **315** | 1.26 | 50.19 | 5.79 | 86.56 | 1.99 | 1.73 |
| **All** | 1.85 | 49.16 | 2.46 | 87.32 | 1.82 | 1.59 |
| **Theta waves in human EEG** |  |  |  |  |  |  |
| **0** | -27.55 | 56.33 | -26.6 | 78.66 | 1.63 | 1.74 |
| **45** | 3.02 | 53.48 | 2.74 | 84.40 | 1.98 | 2.26 |
| **90** | -2.51 | 49.95 | -2.38 | 88.51 | 2.31 | 2.93 |
| **135** | -11.94 | 49.20 | -11.96 | 88.19 | 2.46 | 3.08 |
| **180** | -19.22 | 51.25 | -18.24 | 84.36 | 2.01 | 2.42 |
| **225** | 0.15 | 52.84 | -0.60 | 85.65 | 2.07 | 2.43 |
| **270** | -5.63 | 52.35 | -6.25 | 86.84 | 2.20 | 2.63 |
| **315** | -15.52 | 50.49 | -15.90 | 87.10 | 2.32 | 2.79 |
| **All** | -10.25 | 52.86 | -10.35 | 85.09 | 2.09 | 2.53 |
| **Alpha waves in human EEG** |  |  |  |  |  |  |
| **0** | -18.08 | 55.77 | -17.80 | 81.58 | 1.15 | 1.20 |
| **45** | -24.59 | 55.09 | -21.55 | 79.14 | 0.89 | 0.90 |
| **90** | -13.22 | 54.99 | -12.19 | 84.02 | 0.87 | 0.84 |
| **135** | -7.16 | 56.20 | -6.29 | 83.29 | 1.15 | 1.19 |
| **180** | -24.12 | 56.76 | -23.49 | 79.72 | 1.12 | 1.16 |
| **225** | -27.33 | 55.93 | -25.10 | 78.55 | 0.86 | 0.87 |
| **270** | -14.37 | 54.65 | -14.60 | 82.64 | 0.86 | 0.87 |
| **315** | -4.77 | 55.14 | -3.71 | 86.62 | 1.16 | 1.23 |
| **All** | -17.12 | 55.80 | -16.34 | 81.40 | 0.99 | 1.03 |

EEG = electroencephalography. Inter-stimulus-interval is measured in seconds. Negative phase offset values indicate stimulus presentation before the targeted phase, positive phase offset values indicate stimulus presentation after the targeted phase.

**Table 2.** Average phase-targeting performance for each participant across all targeted phases (0°, 45°, 90°, 135°, 180°, 225°, 270°, and 315°), for slow oscillations, theta and alpha waves in human EEG, and theta waves in rodent LFP.

| Participant | Phase offset (°) |  |  | 180° range (%) | Inter-stimulus-interval (s) |  |
| --- | --- | --- | --- | --- | --- | --- |
|  | Mean | SD | Median | Percentage | Mean | SD |
| **Slow oscillations in human EEG** |  |  |  |  |  |  |
| P1 | -10.7 | 56.32 | -10.71 | 80.89 | 2.54 | 2.33 |
| P2 | 2.60 | 45.48 | 3.82 | 89.85 | 1.60 | 1.09 |
| P3 | -1.43 | 52.41 | -1.38 | 85.37 | 2.09 | 1.98 |
| P4 | 3.15 | 50.51 | 2.92 | 86.15 | 1.75 | 1.46 |
| P5 | 7.13 | 42.91 | 7.36 | 91.24 | 1.44 | 1.07 |
| P6 | 3.55 | 48.49 | 2.92 | 88.03 | 1.74 | 1.35 |
| P7 | -0.07 | 50.58 | 0.27 | 86.17 | 1.97 | 1.72 |
| **Theta waves in human EEG** |  |  |  |  |  |  |
| P1 | -18.34 | 51.35 | -18.07 | 85.33 | 2.78 | 2.94 |
| P2 | -10.25 | 50.90 | -9.02 | 86.77 | 1.53 | 1.67 |
| P3 | -4.13 | 52.37 | -4.90 | 85.90 | 1.57 | 1.87 |
| P4 | -8.29 | 54.26 | -8.87 | 83.95 | 2.51 | 3.12 |
| P5 | -12.59 | 54.05 | -12.3 | 83.78 | 2.68 | 3.24 |
| P6 | -12.36 | 54.67 | -12.08 | 83.30 | 2.26 | 2.47 |
| **Alpha waves in human EEG** |  |  |  |  |  |  |
| P1 | -20.76 | 56.11 | -19.35 | 80.36 | 1.12 | 1.24 |
| P2 | -11.91 | 58.86 | -12.36 | 79.10 | 1.12 | 1.14 |
| P3 | -17.34 | 58.28 | -15.79 | 78.86 | 0.98 | 0.97 |
| P4 | -17.84 | 52.52 | -16.73 | 84.24 | 0.78 | 0.69 |
| P5 | -12.82 | 55.02 | -12.63 | 83.16 | 1.07 | 1.11 |
| **Theta waves in rodent LFP** |  |  |  |  |  |  |
| P1 | -2.76 | 26.33 | -3.04 | 99.25 | 1.03 | 0.73 |
| P2 | -1.72 | 33.08 | -2.70 | 97.21 | 0.85 | 0.60 |
| P3 | -2.13 | 30.41 | -2.74 | 97.91 | 0.89 | 0.62 |

EEG = electroencephalography. LFP = local field potentials. Inter-stimulus-interval is measured in seconds. Negative phase offset values indicate stimulus presentation before the targeted phase, positive phase offset values indicate stimulus presentation after the targeted phase.

Considering accuracy in individual participants (N = 7; figure 2c; supplementary figure 2), the lowest and highest overall marker offsets were −0.07° and −10.70°, while precisions (SD) ranged from 42.91° to 56.32°. The inter-stimulus-interval ranged between 1.44 ± 1.07 and 2.54 ± 2.33 seconds. Together, the findings show that slow oscillation phases can be targeted by M-CLNS with high accuracy, precision, and density. Furthermore, despite minor differences in performance, phase targeting succeeded in all individual participants with high accuracy, precision, and frequency.

### 3.3. Theta waves during REM sleep in human EEG

For theta waves in human EEG, grand average targeting performance across 27074 recorded stimulus onset markers, covering all target phases and participants, was −10.25° ± 52.86°. Of these markers, 85.09% were within a 180° range centered around the target phase (figure 3a). The inter-stimulus-interval, averaged across all markers, was2.09 ± 2.53 seconds.

**Figure 3.**
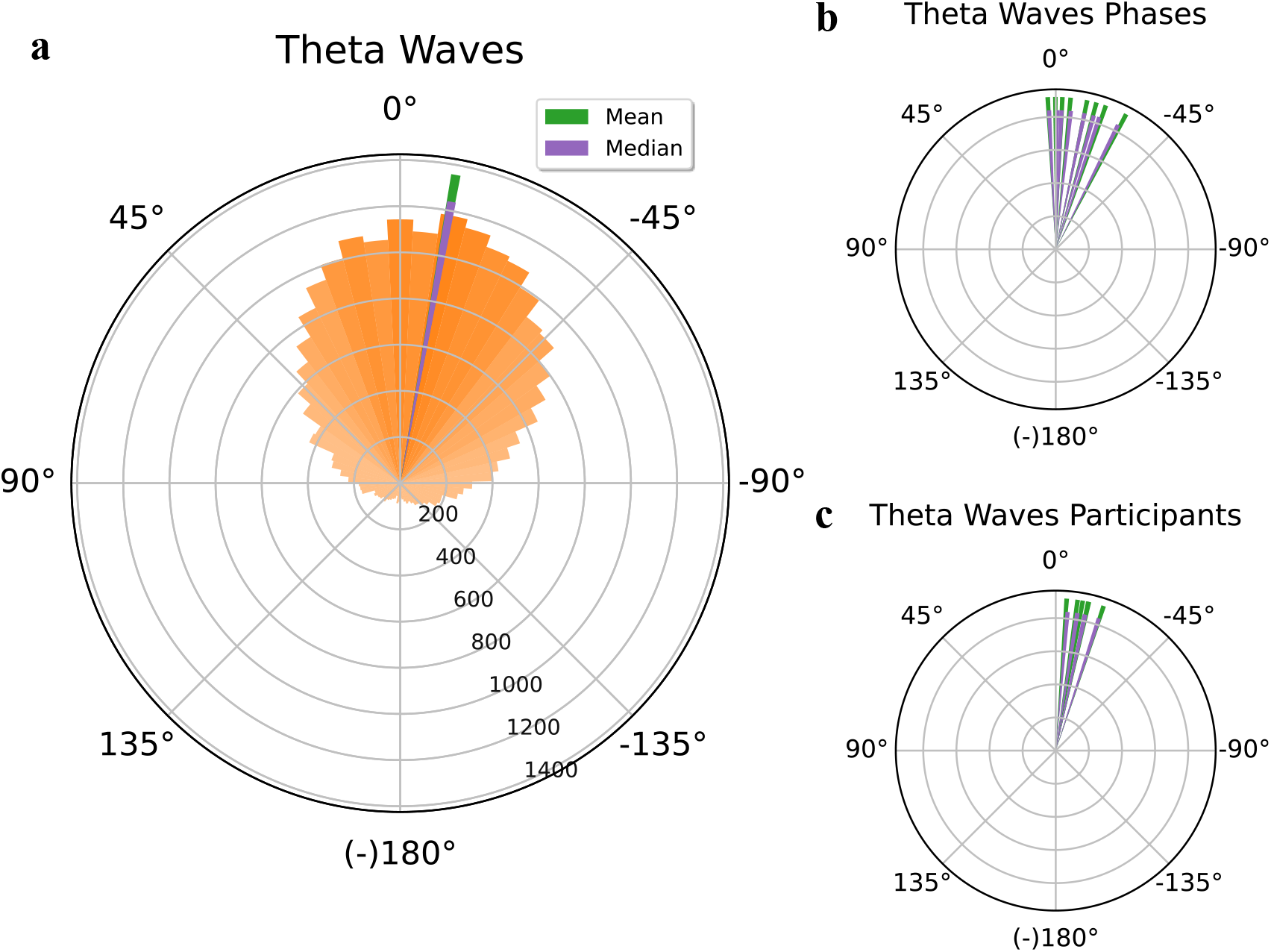
Phase-targeting performance for theta waves in human EEG. (a) Circular histogram showing the distribution of binned stimulus marker phase offsets, defined as the difference between targeted phase and the actual stimulus marker phase, measured in degrees. The circular histogram combines data from all eight targeted phases (0°, 45°, 90°, 135°, 180°, 225°, 270°, and 315°) and all participants (orange) together with the mean (green line) and median (purple line) stimulus marker phase offset. (b) Circular histogram depicting the mean (green lines) and median (purple lines) stimulus marker phase offset for each of the eight individual targeted phases. (c) Circular histogram depicting the mean (green lines) and median (purple lines) stimulus marker phase offset for each individual participant.

Considering performance measures for individual target phases (table 1 and figure 3b; see supplementary figure 3 for polar plots for each targeted phase), offsets ranged between 0.15° ± 52.84° and −27.55° ± 56.33°. The inter-stimulus-interval in seconds ranged between 1.63 ± 1.74 and 2.46 ± 3.08.

In individual participants (N = 6; figure 3c; supplementary figure 4), the lowest and highest offsets were −4.13° and −18.34°, while precisions (SD) ranged from 50.90° to 54.67°, and inter-stimulus-intervals ranged between 1.53 ± 1.67 and 2.78 ± 2.94 seconds.

These findings show that phases around the 360° radial extent of theta waves could be targeted accurately, precisely, and frequently using M-CLNS, in all participants. Compared to targeting in slow oscillations in human EEG, overall accuracy, precision, and density appear slightly lower.

### 3.4. Alpha waves during wake after sleep onset in human EEG

Grand average targeting performance for alpha waves in human EEG, assessed across 40333 stimulus onset markers including all target phases and participants, was −17.12° ± 55.80°. Of these markers, 81.40% were within a 180° range centered around the target phase (figure 4a). The inter-stimulus-interval, averaged across all markers, was0.99 ± 1.03 seconds.

**Figure 4.**
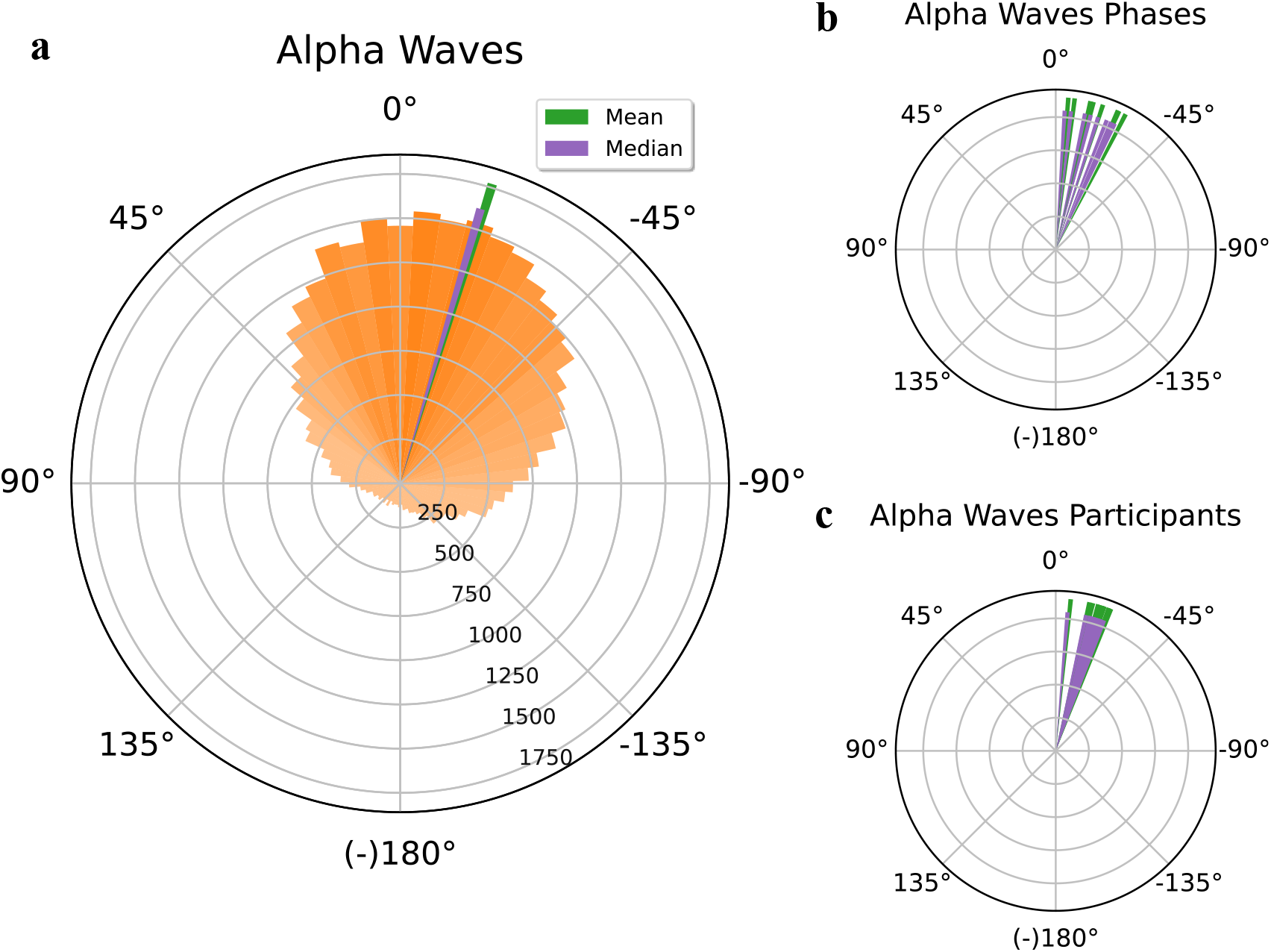
Phase-targeting performance for alpha waves in human EEG. (a) Circular histogram showing the distribution of binned stimulus marker phase offsets, defined as the difference between targeted phase and the actual stimulus marker phase, measured in degrees. The circular histogram combines data from all eight targeted phases (0°, 45°, 90°, 135°, 180°, 225°, 270°, and 315°) and all participants (orange) together with the mean (green line) and median (purple line) stimulus marker phase offset. (b) Circular histogram depicting the mean (green lines) and median (purple lines) stimulus marker phase offset for each of the eight individual targeted phases. (c) Circular histogram depicting the mean (green lines) and median (purple lines) stimulus marker phase offset for each individual participant

Performance measures for individual target phases (table 1 and figure 4b; see supplementary figure 5 for polar plots for each targeted phase) ranged between −4.77° ± 55.14° and −27.33° ± 55.93°, while inter-stimulus-intervals were between 0.86 ± 0.87 and 1.16 ± 1.23.

In individual participants (N = 5), the offsets ranged between −11.91° and −20.76°, while precisions (SD) ranged from 52.52° to 58.86° (figure 4c; supplementary figure 6). The lowest and highest inter-stimulus-intervals were 0.78 ± 0.69 and 1.12 ± 1.24 seconds.

The findings indicate that, overall, alpha wave phases can be predicted by M-CLNS fairly accurately, precisely, and at a high rate, in all participants. Accuracy and precision appear to be slightly lower compared to targeting slow oscillations and theta waves in human EEG, while stimulus density is slightly higher.

### 3.5. Theta waves during REM sleep in rodent hippocampal LFP

For hippocampally recorded theta waves in rodents, the grand average targeting performance across 20292 stimulus onset markers was −2.31° ± 29.47°. Of these markers, 98.31% were recorded within a 180° range centered around the target phase (figure 5a). The mean inter-stimulus-interval across all markers was 0.94 ± 0.67 seconds.

**Figure 5.**
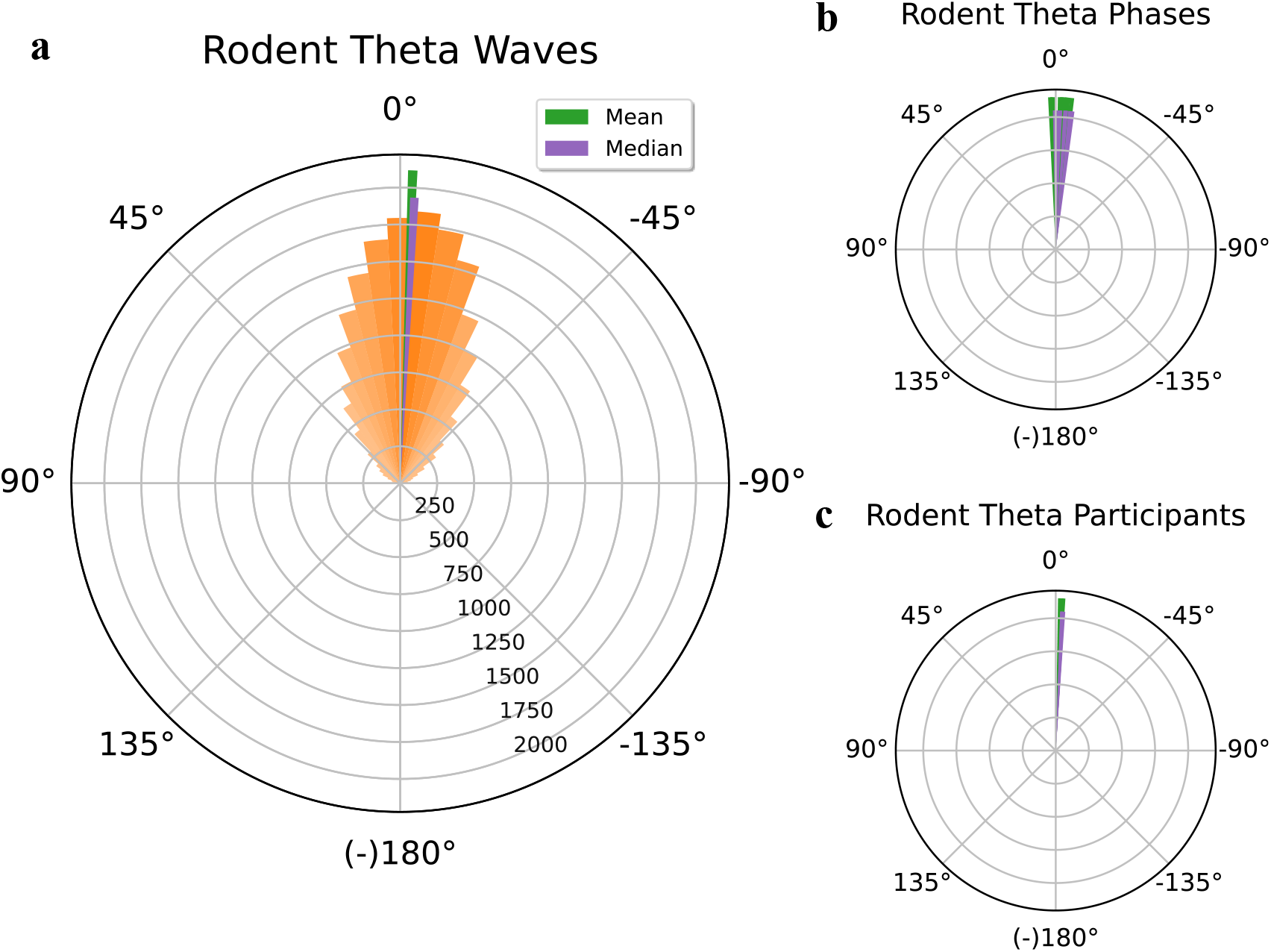
Phase-targeting performance for theta waves in rodent LFP. (a) Circular histogram showing the distribution of binned stimulus marker phase offsets, defined as the difference between targeted phase and the actual stimulus marker phase, measured in degrees. The circular histogram combines data from all eight targeted phases (0°, 45°, 90°, 135°, 180°, 225°, 270°, and 315°) and all participants (orange) together with the mean (green line) and median (purpleline) stimulus marker phase offset. (b) Circular histogram depicting the mean (green lines) and median (purple lines) stimulus marker phase offset for each of the eight individual targeted phases. (c) Circular histogram depicting the mean (green lines) and median (purple lines) stimulus marker phase offset for each individual participant.

Performance for individual target phases (table 3 and figure 5b; see supplementary figure 7 for polar plots for each targeted phase) was very high, with the lowest offset at 1.20° ± 27.31° and the highest at −6.10° ± 28.53°. The inter-stimulus-interval ranged between 0.92 ± 0.66 and 0.95 ± 0.69 seconds.

**Table 3.** Phase-targeting performance for individual and across combined targeted phases (0°, 45°, 90°, 135°, 180°, 225°, 270°, and 315°), for theta waves in rodent LFP.

| Target phase | Phase offset (°) |  |  | 180° range (%) | Inter-stimulus-interval (s) |  |
| --- | --- | --- | --- | --- | --- | --- |
|  | Mean | SD | Median | Percentage | Mean | SD |
| **Theta waves in rodent LFP** |  |  |  |  |  |  |
| **0** | -4.24 | 31.52 | -4.21 | 98.75 | 0.94 | 0.68 |
| **45** | -3.07 | 30.84 | -1.38 | 97.66 | 0.94 | 0.67 |
| **90** | 1.20 | 27.31 | 0.13 | 98.45 | 0.95 | 0.69 |
| **135** | 2.02 | 27.82 | 0.29 | 98.43 | 0.92 | 0.66 |
| **180** | -1.67 | 27.30 | -1.81 | 99.10 | 0.93 | 0.65 |
| **225** | -6.10 | 28.53 | -6.86 | 98.12 | 0.94 | 0.65 |
| **270** | -4.54 | 30.01 | -6.04 | 97.76 | 0.95 | 0.67 |
| **315** | -2.44 | 31.25 | -3.99 | 98.16 | 0.94 | 0.68 |
| **All** | -2.31 | 29.47 | -2.85 | 98.31 | 0.94 | 0.67 |

LFP = local field potentials. Inter-stimulus-interval is measured in seconds. Negative phase offset values indicate stimulus presentation before the targeted phase, positive phase offset values indicate stimulus presentation after the targeted phase.

For individual rodents (N = 3), the lowest and highest offsets were −1.72° and −2.76°, while precisions (SD) ranged from 26.33° to 33.08° (figure 5c; supplementary figure 8). The inter-stimulus-interval ranged between 0.85 ± 0.60 and 1.03 ± 0.73 seconds. These findings show very high performance across all targeting measures; in fact, performance appears superior compared to targeting of any of the human EEG oscillations. Also, performance seems to vary less with target phase compared to in human EEG.

## 4. Discussion

### 4.1. M-CLNS models, tracks, and predicts varied oscillatory dynamics adequately

The aim of this study was to evaluate oscillatory phase targeting performance of M-CLNS, a new closed-loop neurostimulation approach, based on signal modelling. To assess the method’s generic potential, performance was evaluated across different types of neurophysiological signals (human EEG, rodent LFP), oscillatory frequency bands (slow oscillations, theta, alpha) and target phases (8 radial phases equidistantly spread across the 360° range). We observed accurate performance of the method in all tested conditions. Furthermore, performance was adequate for all individual participants at each hallmark oscillation and phase. While performance varied across target phases, for all hallmark oscillations, these variations did not show any consistency with respect to the more and less accurately targeted phases. This is, to our knowledge, the first and only method for which such flexibility has been demonstrated.

While phase targeting performance was adequate across all tested hallmark oscillations, it varied somewhat with targeted frequency band. Several factors may underly these variations: first, the frequency of the neural oscillation plays a role, with phase targeting of faster dynamics being progressively more demanding. Note that 1^o^ of a 1 Hz (slow oscillation range) and a 10 Hz wave (alpha wave range) correspond to 2.8 ms and 0.28 ms, respectively. Thus, phase targeting of faster dynamics requires progressively higher temporal precision of the closed-loop system. This circumstance might underly the progressively lower precision we observed in phase targeting slow oscillations, theta and alpha waves in the human EEG. Another likely factor influencing performance regards the phase regularity of the targeted oscillatory dynamic.

The more regular the dynamic, the more accurate the phase predictions will tend to be, while phase jitter from one oscillation to the next will tend to induce phase targeting error. Across the hallmark oscillations, the highest phase targeting performance by far was achieved targeting theta waves in rodent LFPs. The probable reason is that, compared to human scalp EEG recordings, this depth recording presents a more (phase) regular signal. Contributing factors hereto may include the higher signal to noise ratio and smaller signal source field of LFPs compared to EEG, as well as inherent regularity of hippocampal theta in rodents.

The high performance and broad applicability of the M-CLNS approach is based on a signal modelling procedure that is, in principle, applicable to any oscillatory signal, and provides a prediction of signal continuation. As demonstrated, the prediction allows for stimuli to be precisely targeted to future time points where a particular oscillatory phase is expected to occur.

In contrast to the flexibility of M-CLNS, most other reported CLNS procedures target one specific pattern of interest in a signal and, as such, have narrow applicability. Moreover, in such approaches, a stimulus is released upon target pattern detection, meaning that the stimulus is delayed relative to the pattern of interest, as a function of the lag in the closed-loop system.

A further advantage of M-CLNS, lies in the lack of real-time signal filtering. Such fast filtering, which is a necessary aspect of other reported CLNS procedures, induces additional looplag, as well as signal distortion, both of which negatively affect targeting performance. A final benefit of M-CLNS over previously reported methods [9,17,22], is that due to the intrinsic adaptability to the input signal there is no need for individual calibration sessions or training on large data sets representative of specific target populations. The elimination of such time-consuming procedures may favor the broad applicability of M-CLNS.

In line with the intended generic applicability of our approach, the method has a software-based implementation. The software features a graphical user-interface that allows adaptation of all relevant algorithm parameters, including size of the calculus window (stretch of most recent data on which the real-time analysis is executed) and search window, target frequency band, target phase, model fit threshold, model amplitude threshold (optional), and the size of any imposed inter-stimulus-interval (optional). Through these modifiable parameter settings, the procedure can be conveniently adapted and optimized to new applications. Examples of such applications may include targeting other EEG frequency bands during wakefulness or sleep, or using other types of oscillatory, electrophysical signals, such as ECG (electrocardiogram), or human intracranial EEG.

### 4.2. Technical considerations

An important consideration in dealing with CLNS procedures concerns loop lag, that is, the processing time of the algorithm plus the throughput time of all other hardware and software components in the closed loop. While our modelling approach provides a signal prediction that is used to compensate for loop lag, minimizing loop lag is still essential, as the model prediction must still be valid beyond the loop lag. Given the high time variance of neurophysiological signals, this means that any time loss may be deleterious to performance. Thus, all hardware and software components of the closed loop should have short throughput times with minimal jitter. Given the crucial importance of speed, the modelling procedure executed by the algorithm was developed to be highly time efficient, with the sole computations at each calculation step being the sine fitting and the calculation of a simple measure of model fit.

As a further consideration, while phase-targeting performance of M-CLNS as shown in our results is adequate for many applications, any systematic offsets from a given target can be corrected by simply adding or subtracting the mean temporal error as a fixed variable in the algorithm, thus further enhancing targeting accuracy. Such adjustments were, however, not applied in this study nor in any of our experimental studies with M-CLNS.

### 4.3. Future perspectives

While the current study only concerns phase targeting performance of M-CLNS, experimental studies provide insight into the effects of phase targeted acoustic stimulation [26-29,41,42]. First of all, a few recent studies indicate that the phase relation of stimuli with intrinsic oscillatory dynamics, and the precision of phase targeting, matter for the effect [39,40]. Furthermore, several studies in healthy human subjects show important effects of slow oscillation phase targeted stimulation on measures of sleep depth and memory consolidation [8,9], while theta phase targeted stimulation influences emotional memory consolidation and dream content during REM sleep [29]. Additionally, emerging evidence in clinical populations, such as those with PTSD, suggests that M-CLNS holds promise for therapeutic applications [41,42]. Indeed, many sleep disorders, as well as healthy aging, are associated with a reduction in slow oscillation amplitude and overall sleep depth. Closed-loop slow oscillation boosting is being explored as a novel, non-invasive approach to enhance sleep quality [15,16,24] Non-adaptive methods may struggle to target the shallow slow oscillation dynamics prevalent in elderly and clinical populations, particularly because these methods rely on amplitude thresholds.

Conversely, the ability of M-CLNS to accurately target oscillatory phase without requiring an amplitude criterion makes it uniquely suited for populations with compromised slow oscillation dynamics. For instance, in a study involving PTSD patients, M-CLNS successfully targeted slow oscillation phases with high accuracy, using the same algorithm as in healthy young subjects [42]. Furthermore, M-CLNS was used to phase-target posterior slow oscillations, which exhibit much lower amplitudes than frontal ones [28]. Pilot studies in individuals with insomnia have also shown promising results, with high accuracy in slow oscillation phase targeting. These findings indicate significant potential for developing M-CLNS as a therapeutic tool for sleep enhancement.

As a step towards clinical application, we have recently miniaturized M-CLNS into a research-grade wearable sleep stimulation device, designed for home use. This self-applicable device will enable longer-term studies, assessing the effects and underlying mechanisms of phase-targeted acoustic stimulation in natural situations, like the home environment. It should, moreover, facilitate studies on the potential therapeutic effects of M-CLNS across various clinical populations, including those affected by insomnia, PTSD, Alzheimer’s disease, schizophrenia and autism spectrum disorder.

## 5. Conclusion

In conclusion, the demonstrated flexibility of M-CLNS, combined with its real-time functioning without need for calibration, makes it a promising tool for advancing the field of closed-loop neurostimulation. We anticipate that the generic applicability and convenient software-based implementation of M-CLNS will stimulate and facilitate further development of CLNS procedures and applications.

## Data availability statement

The data that support the findings of this study are available upon reasonable request from the authors.

## Acknowledgements

We thank Eva van Poppel (†), Roy Cox, and Ilia Korjoukov for their contributions to our work on CLNS algorithms.

## Conflict of interest

Lucia Talamini is an inventor of the patented modelling-based closed-loop neurostimulation method utilized in this study. This patent may lead to potential financial interests. The authors declare that this does not affect the objectivity of the research findings.

## Author contributions

Conceptualization & Methodology: S L J and L M T; Formal Analysis: S L J; Simulations data collection: S L J; Human subjects data collection: A C V D H and S L J; Animal subjects data collection: J P; Supervision: L M T; Writing: all authors contributed to manuscript writing and editing; All authors have read and agreed to the published version of the manuscript.

## Appendix. Supplementary figures

**Supplementary figure 1.**
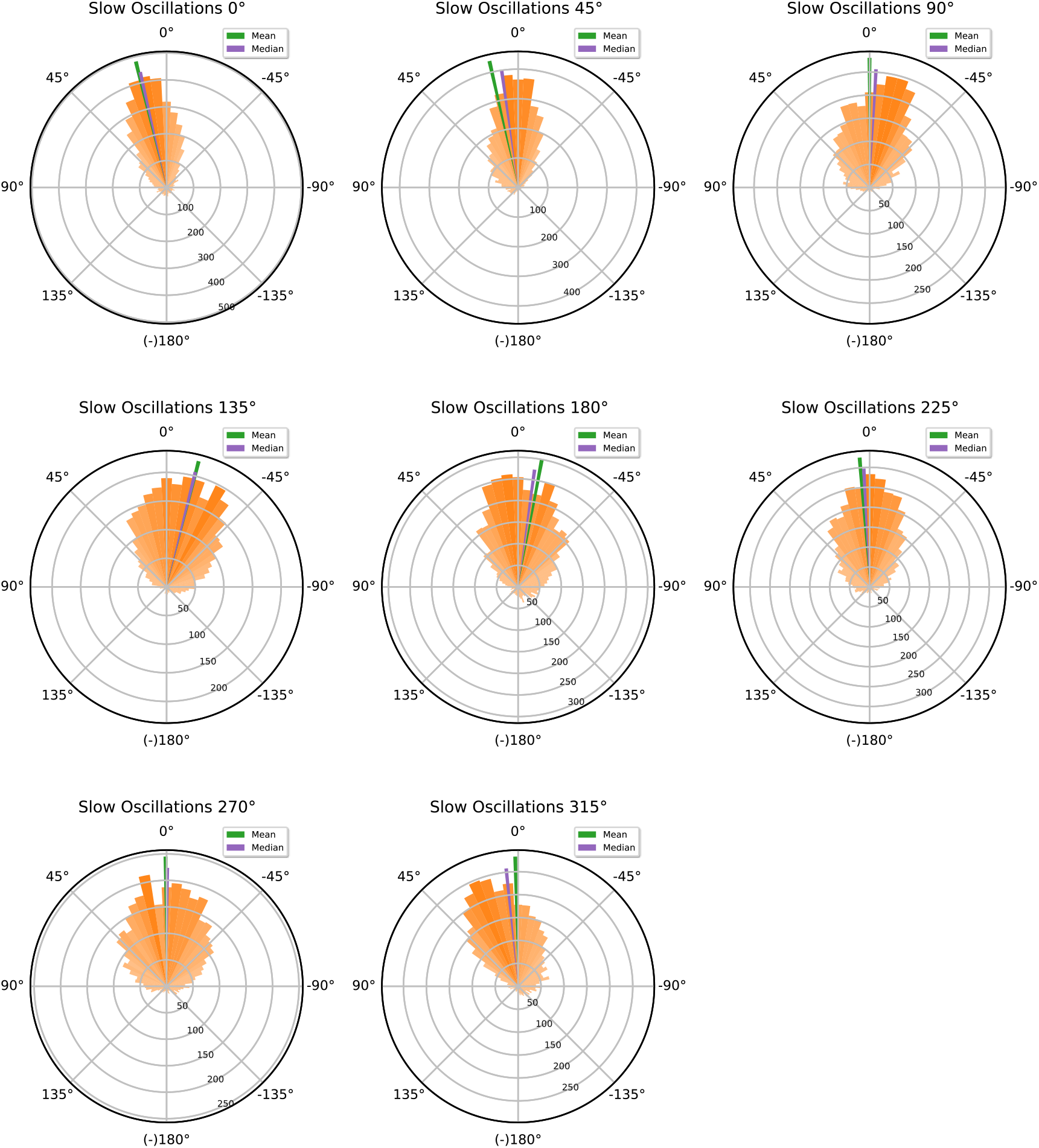
Phase-targeting performance for slow oscillations in human EEG for each targeted phase (0°, 45°, 90°, 135°, 180°, 225°, 270°, and 315°). The circular histograms show the distribution of binned stimulus marker phase offsets across all participants (orange), defined as the difference between each stimulus marker phase and the targeted phase, measured in degrees, together with the mean (green line) and median (purple line) stimulus marker phase offset.

**Supplementary figure 2.**
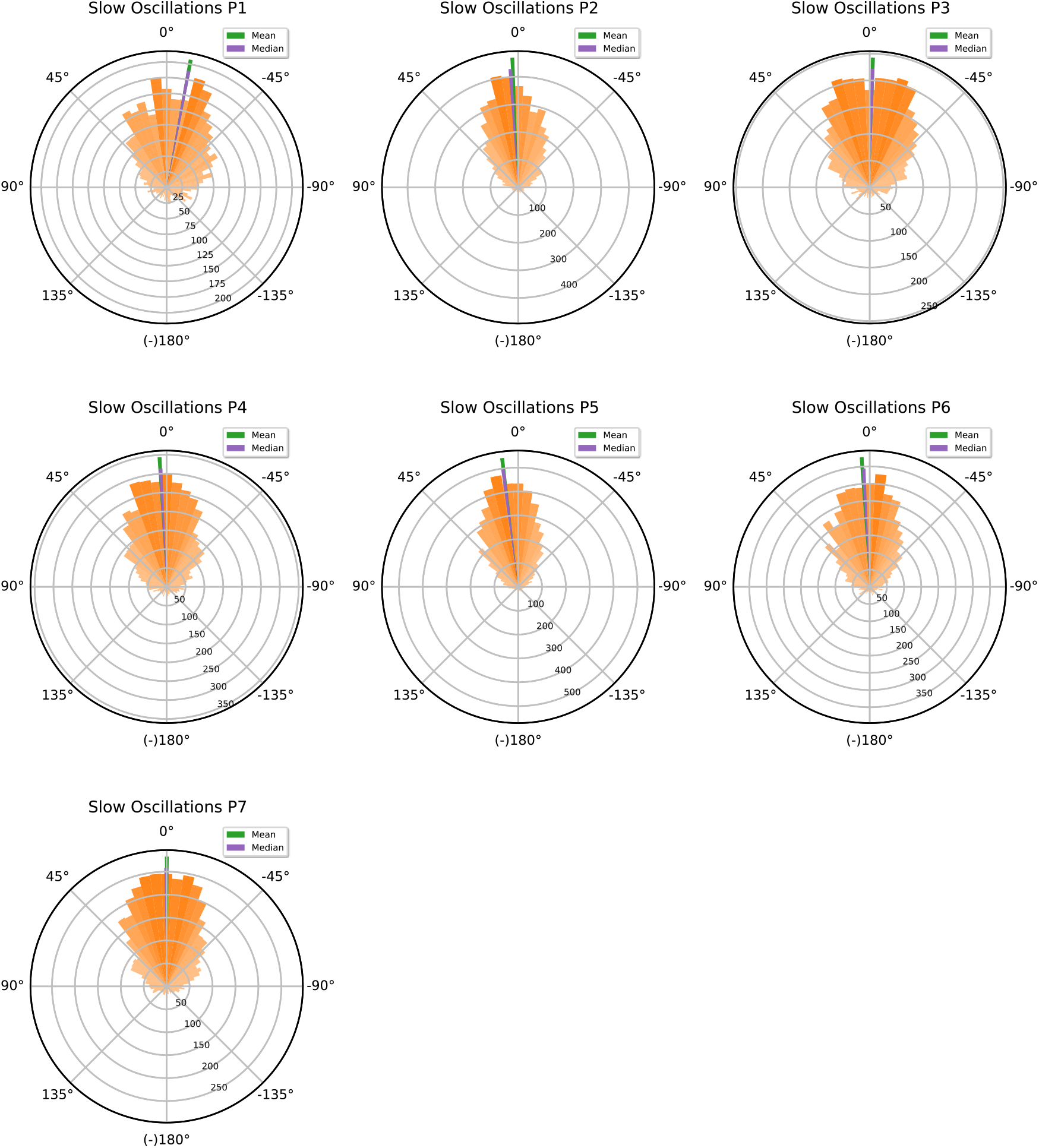
Phase-targeting performance for slow oscillations in human EEG for each participant across all eight targeted phases (0°, 45°, 90°, 135°, 180°, 225°, 270°, and 315°). The circular histograms show the distribution of binned stimulus marker phase offsets across all eight phases (orange), defined as the difference between each stimulus marker phase and the targeted phase, measured in degrees, together with the mean (green line) and median (purple line) stimulus marker phase offset.

**Supplementary figure 3.**
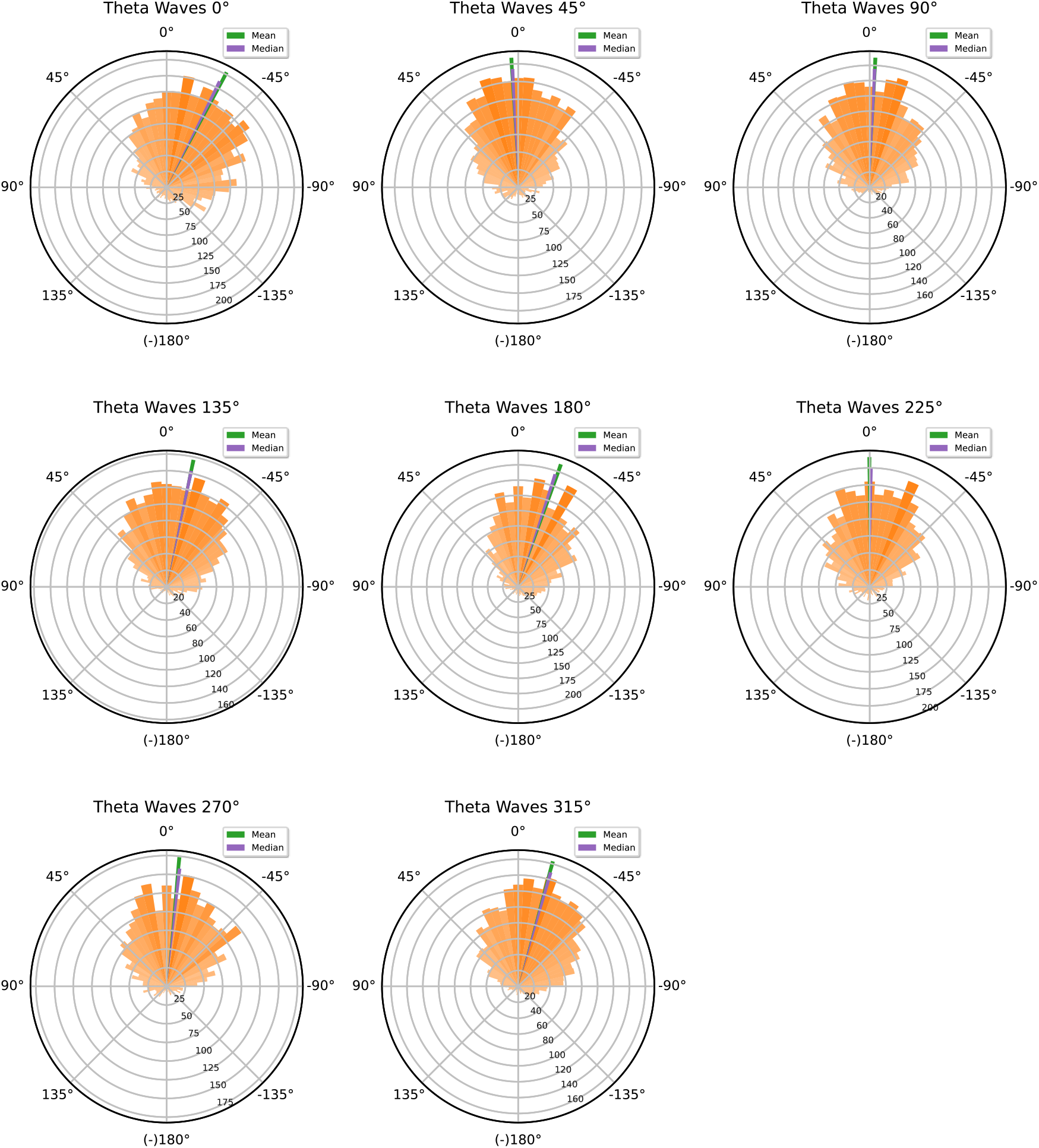
Phase-targeting performance for theta waves in human EEG for each targeted phase (0°, 45°, 90°, 135°, 180°, 225°, 270°, and 315°). The circular histograms show the distribution of binned stimulus marker phase offsets across all participants (orange), defined as the difference between each stimulus marker phase and the targeted phase, measured in degrees, together with the mean (green line) and median (purple line) stimulus marker phase offset.

**Supplementary figure 4.**
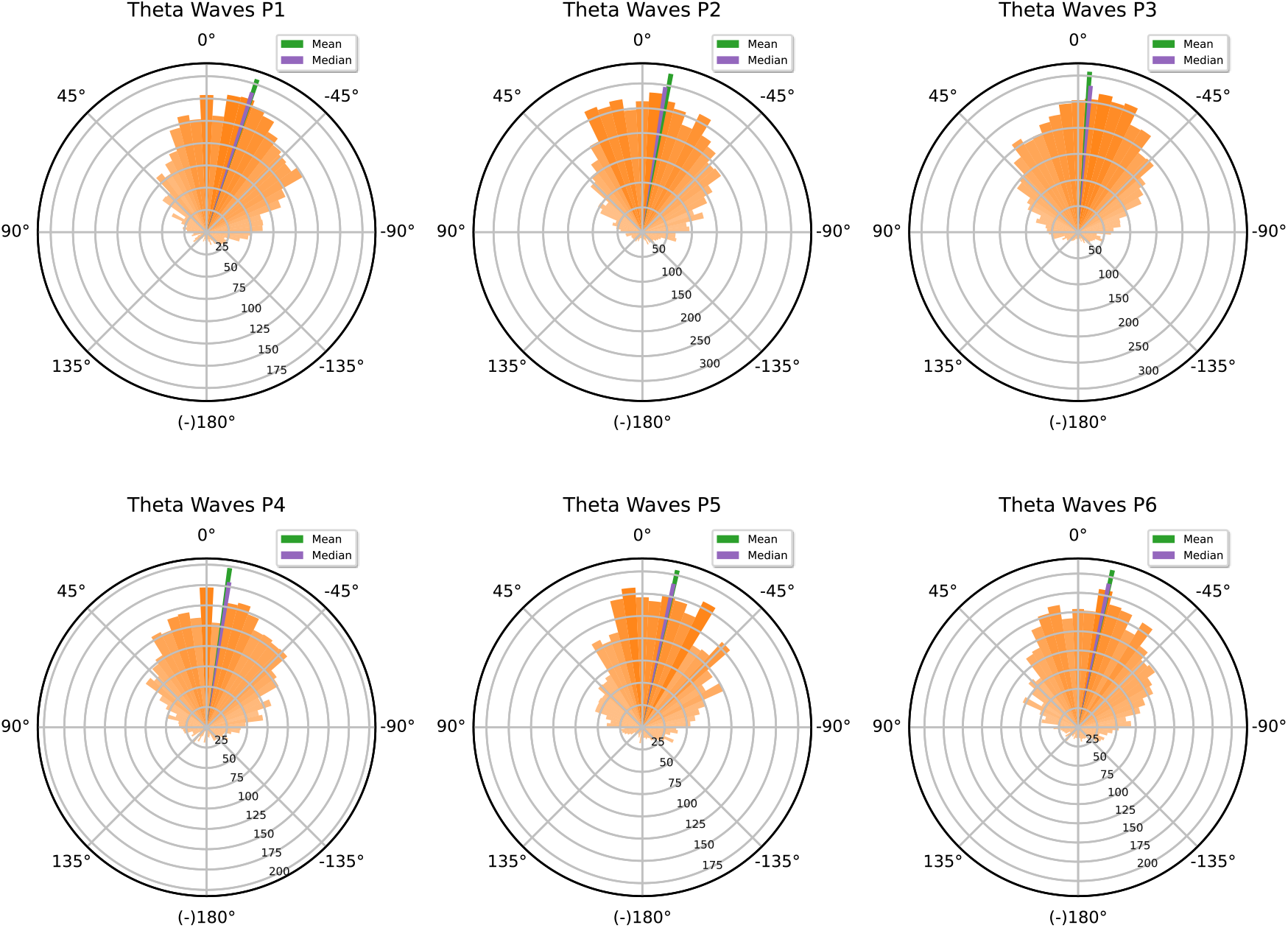
Phase-targeting performance for theta waves in human EEG for each participant across all eight targeted phases (0°, 45°, 90°, 135°, 180°, 225°, 270°, and 315°). The circular histograms show the distribution of binned stimulus marker phase offsets across all eight phases (orange), defined as the difference between each stimulus marker phase and the targeted phase, measured in degrees, together with the mean (green line) and median (purple line) stimulus marker phase offset.

**Supplementary figure 5.**
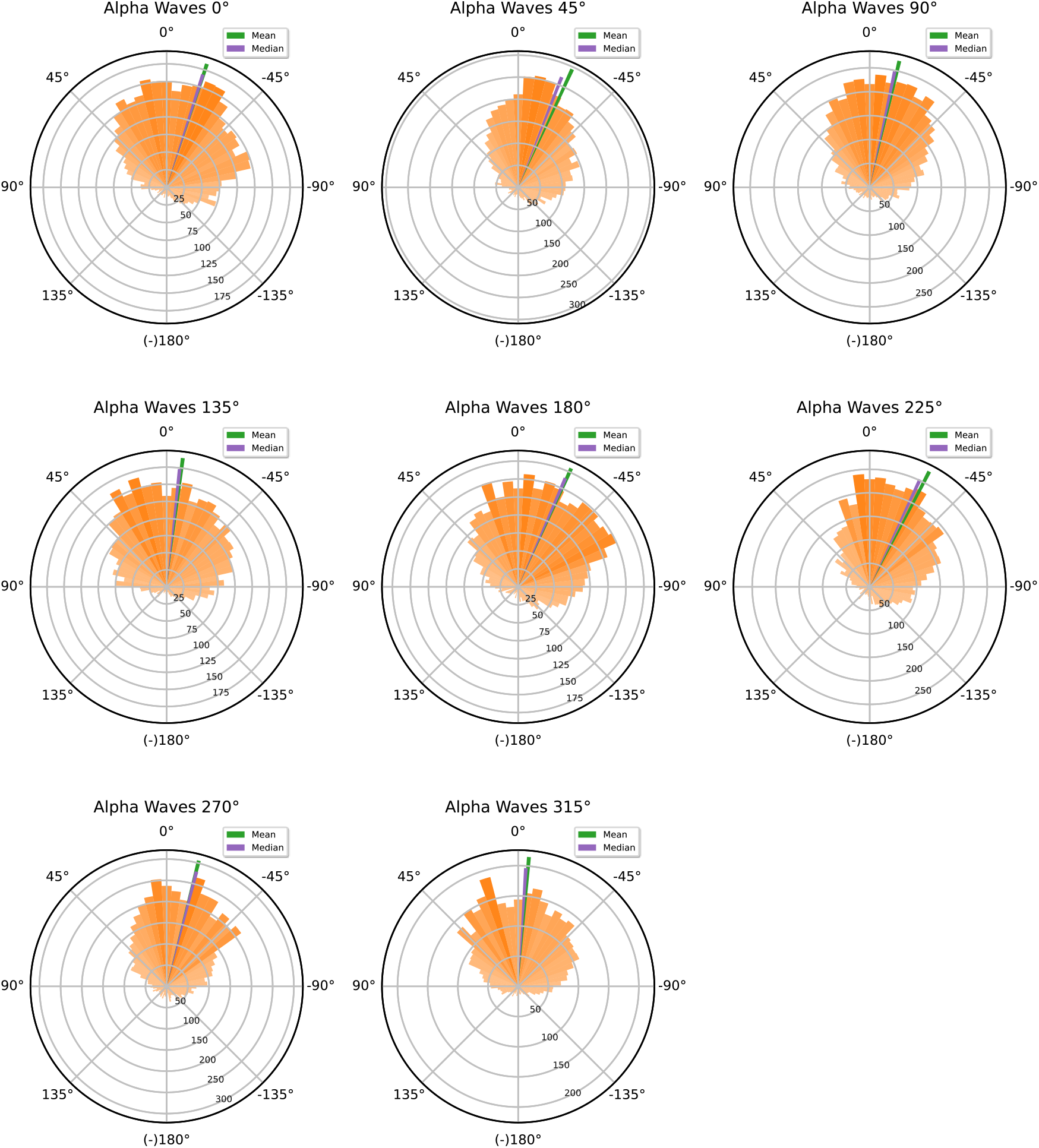
Phase-targeting performance for alpha waves in human EEG for each targeted phase (0°, 45°, 90°, 135°, 180°, 225°, 270°, and 315°). The circular histograms show the distribution of binned stimulus marker phase offsets across all participants (orange), defined as the difference between each stimulus marker phase and the targeted phase, measured in degrees, together with the mean (green line) and median (purple line) stimulus marker phase offset.

**Supplementary figure 6.**
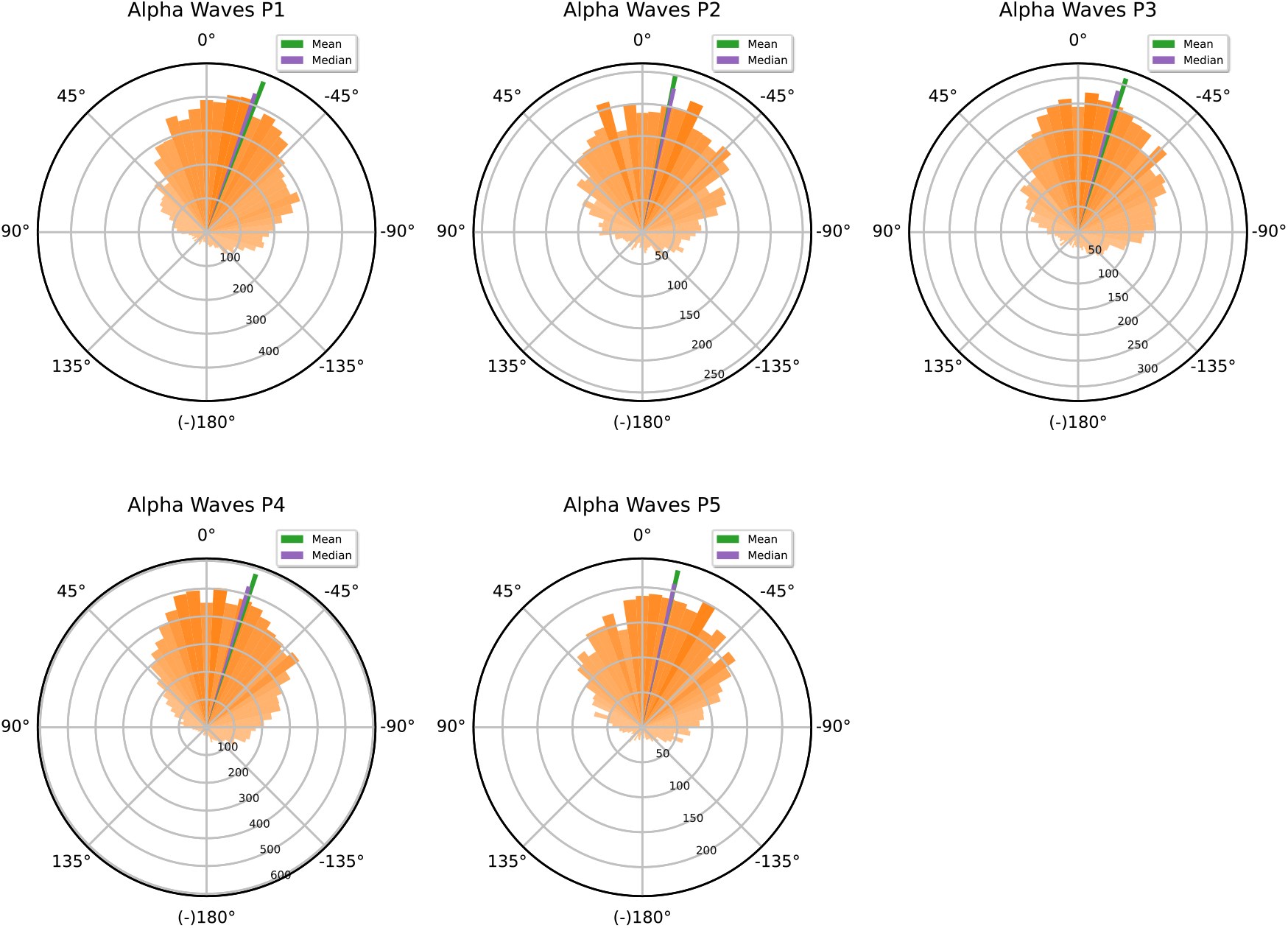
Phase-targeting performance for alpha waves in human EEG for each participant across all eight targeted phases (0°, 45°, 90°, 135°, 180°, 225°, 270°, and 315°). The circular histograms show the distribution of binned stimulus marker phase offsets across all eight phases (orange), defined as the difference between each stimulus marker phase and the targeted phase, measured in degrees, together with the mean (green line) and median (purple line) stimulus marker phase offset.

**Supplementary figure 7.**
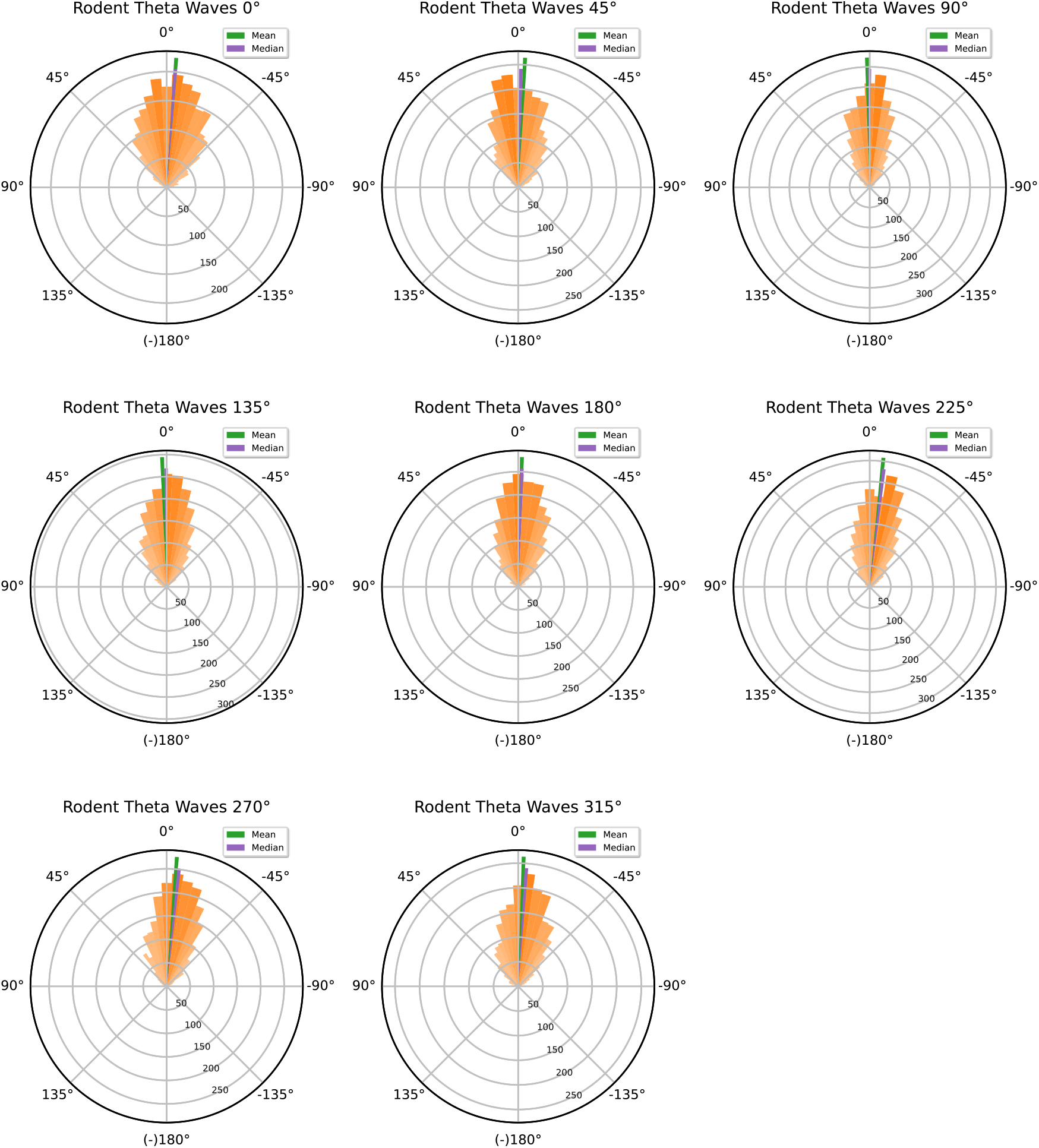
Phase-targeting performance for theta waves in rodent LFP for each targeted phase (0°, 45°, 90°, 135°, 180°, 225°, 270°, and 315°). The circular histograms show the distribution of binned stimulus marker phase offsets across all participants (orange), defined as the difference between each stimulus marker phase and the targeted phase, measured in degrees, together with the mean (green line) and median (purple line) stimulus marker phase offset.

**Supplementary figure 8.**
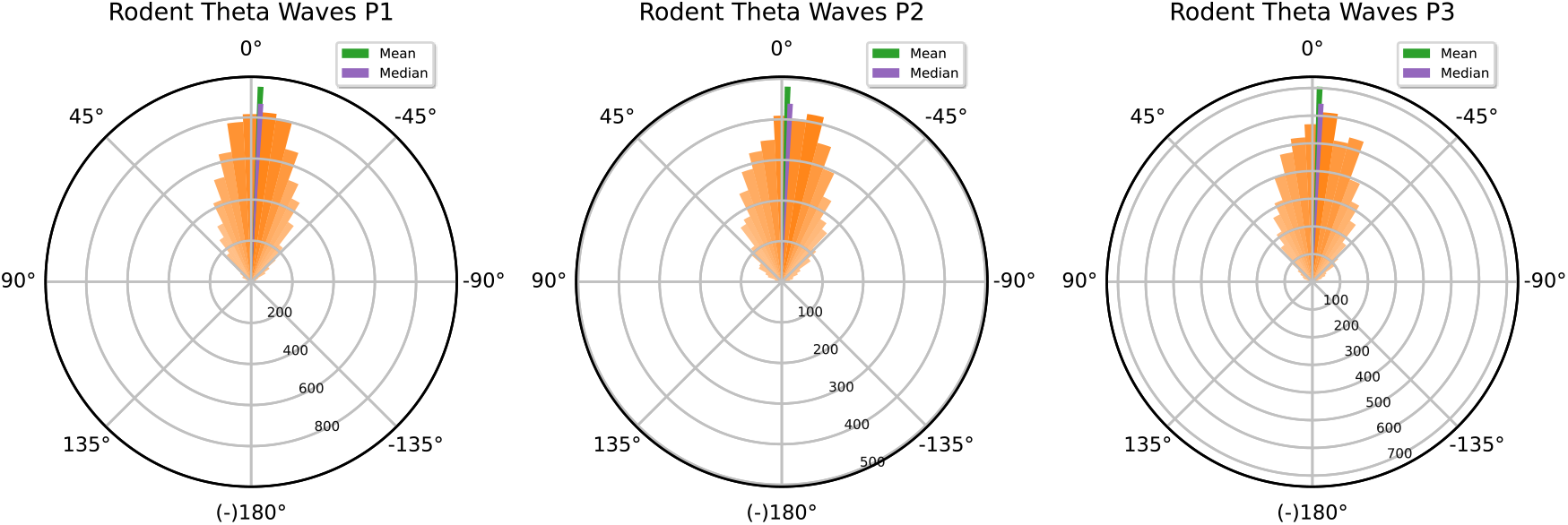
Phase-targeting performance for theta waves in rodent LFP for each participant across all eight targeted phases (0°, 45°, 90°, 135°, 180°, 225°, 270°, and 315°). The circular histograms show the distribution of binned stimulus marker phase offsets across all eight phases (orange), defined as the difference between each stimulus marker phase and the targeted phase, measured in degrees, together with the mean (green line) and median (purple line) stimulus marker phase offset.

## Notes

### Summary of Updates

This version of the manuscript has been revised to update the text of the introduction and the discussion sections for clarity, brevity and balance across the discussed background literature.

